# Widespread regulation of the maternal transcriptome by Nanos in Drosophila

**DOI:** 10.1101/2023.08.28.555109

**Authors:** Mohammad Marhabaie, Tammy H. Wharton, Sung Yun Kim, Robin P. Wharton

**Affiliations:** Nationwide Children’s Hospital; Ohio State University

**Author notes:** Current address: The Steve and Cindy Rasmussen Institute for Genomic Medicine Abigail Wexner Research Institute at Nationwide Children’s Hospital Columbus, OH 43215. Corresponding author: Department of Molecular Genetics, Department of Cancer Biology and Genetics Center for RNA Biology, Ohio State University 484 W. 12th Avenue Columbus, OH 43210.

## Abstract

The translational repressor Nanos (Nos) regulates a single target, maternal *hunchback* (*hb)* mRNA, to govern abdominal segmentation in the early Drosophila embryo. Nos is recruited specifically to sites in the 3’-UTR of *hb* mRNA in collaboration with the sequence-specific RNA-binding protein Pumilio (Pum); on its own, Nos has no binding specificity. Nos is expressed at other stages of development, but very few mRNA targets that might mediate its action at these stages have been described. Nor has it been clear whether Nos is targeted to other mRNAs in concert with Pum or via other mechanisms. In this report, we identify mRNAs targeted by Nos via two approaches. In the first method, we identify mRNAs depleted upon expression of a chimera bearing Nos fused to the nonsense mediated decay (NMD) factor Upf1. We find that, in addition to *hb*, Upf1-Nos depletes ∼2600 mRNAs from the maternal transcriptome in early embryos. Virtually all of these appear to be targeted in a canonical, *hb*-like manner in concert with Pum. In a second, more conventional approach, we identify mRNAs that are stabilized during the maternal zygotic transition (MZT) in embryos from *nos*^-^ females. Most (86%) of the 1185 mRNAs regulated by Nos are also targeted by Upf1-Nos, validating use of the chimera. Approximately 60% of mRNAs targeted by Upf1-Nos are not stabilized in the absence of Nos. However, Upf1-Nos mRNA targets are hypo-adenylated and inefficiently translated at the ovary-embryo transition, whether or not they suffer Nos-dependent degradation in the embryo. We suggest that the late ovarian burst of Nos represses a large fraction of the maternal transcriptome, priming it for later degradation by other factors during the MZT in the embryo.

## Introduction

Nanos is a conserved cytoplasmic repressor that binds to mRNAs, recruiting effectors that block translation and promote degradation (1,2). A wealth of genetic evidence reveals that Nanos is primarily active in the germline, where it controls various aspects of development in a number of organisms. In Drosophila, mouse, and humans, Nos is required for the maintenance of germline stem cells (3–5). Later in germline development, Nos regulates common pathways in the primordial germ cells (PGCs) of Drosophila (6–9) and C. elegans (10,11), including the repression of somatic gene transcription, inhibition of proliferation, and escape from apoptosis.

Evidence of a role for Nos in somatic cells is more limited. In the prospective somatic cytoplasm, Nos has been thought to have one major regulatory target for specification of abdominal segmentation, maternal *hb* mRNA (discussed below) (12–14). At later stages of development, Nos regulates dendritic arborization in the peripheral nervous system and the structure of the neuro-muscular junction of larvae (15–17). In other somatic tissues, a latent capacity of Nos to regulate gene expression is revealed in the absence of *lethal* (*3*) *malignant brain tumor* [*l*(*3*)*mbt*], which encodes a component of two transcriptional repressor complexes (dREAM and LINT). In *l*(*3*)*mbt* mutant larval brains and somatic ovarian cells, expression of Nos (and other germline-restricted genes) is derepressed; remarkably, although hundreds of transcripts are derepressed in the absence of *l*(*3*)*mbt*, mutant phenotypes in both tissues are completely suppressed by loss of Nos activity (18,19). A similar latent regulatory capacity of Nos has been identified in cultured mammalian cells lacking dREAM activity (20). In Drosophila (and likely other organisms as well), both the expression of Nos in the germline and its repression in somatic cells is important for normal development.

Given the diversity of its biological roles, it is likely that Nos regulates a number of mRNAs in various organisms. Support for such an idea comes from the Seydoux lab, who compared the maternal transcriptomes in wild type and *nos* mutant PGCs from early C. elegans embryos, before the onset of zygotic transcription (10). They found that levels of 171 maternal mRNAs are up-regulated in the absence of the redundant function of both Nos-1 and Nos-2, which are required for normal development and survival of the PGCs. It is unclear how many additional maternal mRNAs are translationally repressed but not destabilized, and therefore undetected in their experiments. Later in embryonic PGC development, 871 PGC mRNAs are up-regulated in the absence of Nos function as a result of both delayed turnover of maternal mRNAs and inappropriate zygotic transcription. Nos is not thought to directly regulate transcription, and so presumably acts indirectly to stimulate accumulation of new, zygotic mRNAs.

The mechanism of Nos-dependent regulation is best understood for repression of maternal *hb* mRNA in Drosophila. On its own, Nos binds with high affinity but no detectable sequence specificity to RNA (21,22); however, in collaboration with Pumilio (Pum), it is recruited specifically to composite binding sites (<u>N</u>anos <u>R</u>esponse <u>E</u>lements, NREs) in the *hb* mRNA 3’-UTR (22,23). The composite sites have the consensus sequence 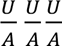 UGUA, with Nos recognizing the first three degenerate positions and Pum the next four. Within the Pum/Nos/RNA ternary complex, amino acid interactions along a protein-protein interface mediate conformational changes in both proteins that enhance binding. A key component of the interface is the Nos C-terminal tail, which is essential for RNA binding; seven amino acids in the tail of the Nos^L7^ mutant protein are absent, and as a result it is not recruited into a ternary complex and does not regulate *hb* mRNA in the embryo (22–24).

Despite an atomic-level understanding of Nos recruitment to the *hb* NRE, two outstanding questions about Nos activity in Drosophila remain. First, how many mRNAs does Drosophila Nos regulate in addition to *hb* (and the handful referenced above)? And second, is Nos recruited to other regulatory targets via its C-terminal tail in collaboration with Pum, as has generally been assumed; or do other RNA-binding proteins recruit Nos via a different mechanism? If Nos bound to a long, information-rich sequence that were rigidly constrained, analysis of the transcriptome sequence might be sufficient to identify likely regulatory targets. But the consensus NRE has low information content and mutations in the NBS portion of the site have only a modest effect on activity in vivo (25). So even though 6225 3’-UTRs in the total Drosophila transcriptome bear one or more consensus NREs (26), it is unclear whether these are genuine regulatory targets in vivo.

To address these issues, we first attempted to identify mRNAs that co-purify with Nos, a method that has been used successfully to identify Pum targets, for example (27,28). Initial reconstruction experiments revealed a disappointingly small enrichment of *hb* mRNA. We therefore turned to an unconventional approach, expressing in flies a protein with the Nonsense Mediated Decay (NMD) factor Upf1 fused to Nos. Early work on NMD showed that tethering Upf1 via an exogenous RBD to a mRNA could direct its degradation (29,30); we reasoned that tethering Upf1 to mRNAs via Nos might lead to their degradation, and that Nos regulatory targets would be depleted from the transcriptome in RNAseq experiments.

## Results

As shown in Figure 1A, Nos is expressed at four important stages of oogenesis and early embryogenesis (24,31,32): (1) in the germline stem cells (GSCs), where it is required for maintenance of stem cell status; (2) in a high-level burst at stage 10B in the nurse cells just prior to a general shutdown of transcription as the egg chamber nears maturation; (3) during oogenesis stages 13-14 and continuing during syncytial nuclear cleavage cycles 1-9 in a gradient emanating from the posterior pole of the embryo, where it represses translation of maternal *hb* mRNA; (4) at nuclear cycle 10 in the PGCs, where it regulates proliferation, blocks apoptosis, and is required for migration into the somatic gonad much later in embryogenesis. After division of the GSC, expression of Nos is turned off in the differentiating daughter cell (the cystoblast) and subsequently re-expressed at various stages of oogenesis. No role has been ascribed to Nos in these stages of oogenesis, including the burst at stage 10B.

**Figure 1.**
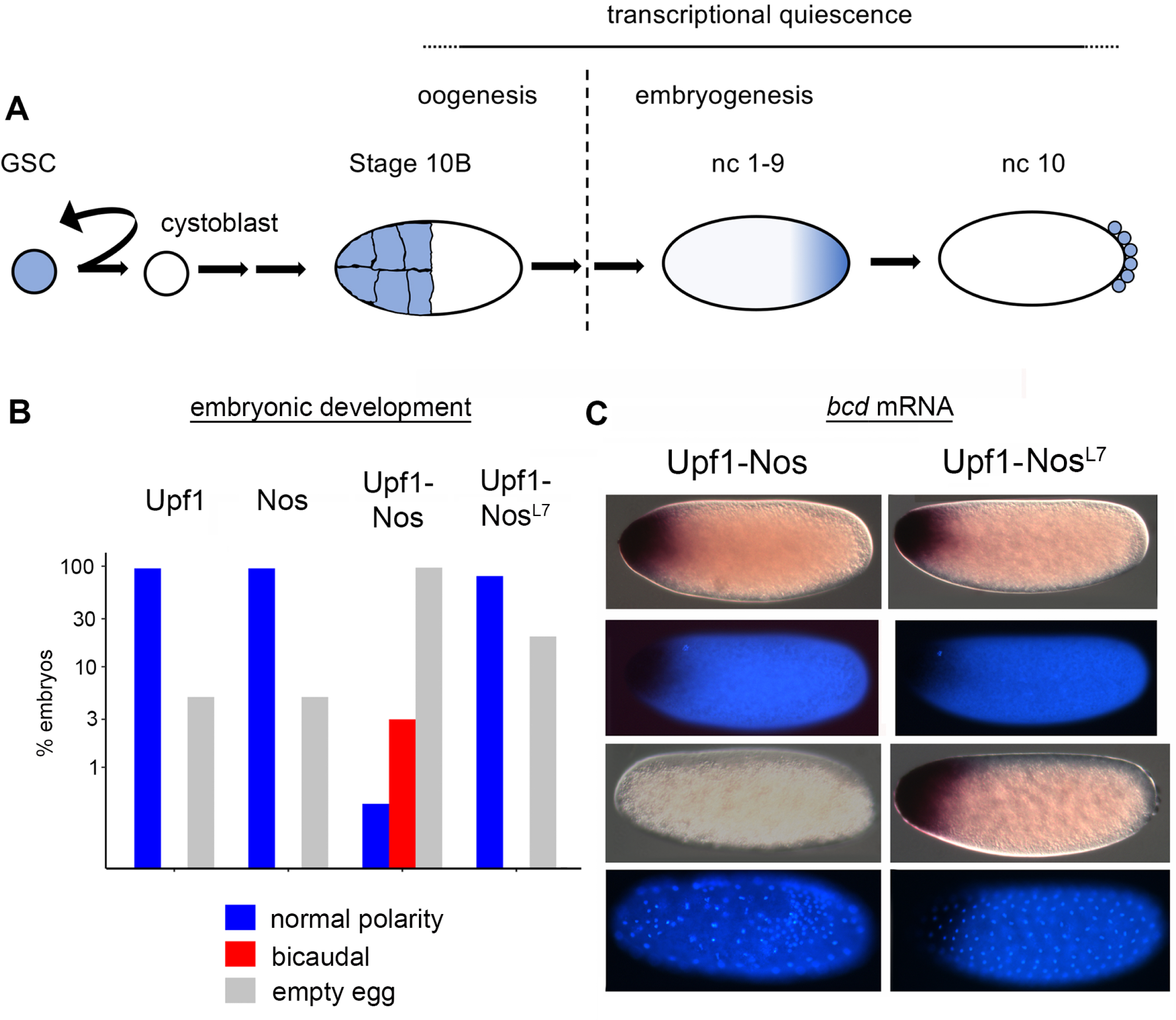
Upf1-Nos suppresses activity of anterior determinants *bicoid* and *hunchback*. A. Timeline of Nos expression during oogenesis and early embryogenesis, focusing on four developmental stages. Note that nuclear cycle (nc) 10 occurs at ∼ 90 minutes of embryonic development. The ∼10-hour period of transcriptional quiescence preceding harvesting of mRNA samples for subsequent analysis is noted above. B. Embryonic phenotypes resulting from maternal expression of the four regulatory proteins, as indicated. For the bargraph (note the logarithmic y-axis), embryos were aged ≥ 24 hours (which allows wild type to hatch) and then scored by sorting into three bins: cuticle with normal polarity (either hatched or not), cuticle with anterior polarity inversion and a bicaudal body plan, or no cuticle (empty egg), which includes both unfertilized eggs and embryos that die prematurely. C. The distribution of *bcd* mRNA in Upf1-Nos and Upf1-Nos^L7^ embryos at two different stages of early embryogenesis--nuclear cycle (nc) 1 in the first two rows and nc 9 in the second two rows. At each stage, two micrographs of a single embryo are shown, using Nomarski optics to visualize *bcd* mRNA and morphology (above), and fluorescence methods to visualize DAPI-stained DNA (below). Statistical analysis is reported in S1 Data. Note that Upf1-Nos embryos appear to be unusually fragile during the fixation for in situ hybridization; many are lost on the wall of the glass tube or at the heptane/aqueous interface.

### Expression of Upf1-Nos inhibits activity of the anterior determinants Hb and Bicoid (Bcd) and causes the degradation of *bcd* mRNA

With the goal of identifying mRNA regulatory targets of Nos, we prepared transgenic flies that express either a Upf1-Nos chimera, or one of three control proteins: Upf1; Nos; or Upf1-Nos^L7^, which bears the C-terminal tail deletion that abrogates recruitment to *hb* mRNA. Transcription of each transgene is dependent on the GAL4 driver that is introduced by appropriate crosses, and each transgenic mRNA bears the 3’-UTR of wildtype *nos* mRNA to provide the post-transcriptional regulation conferred on native *nos*.

We first drove expression of Upf1-Nos and the three control proteins described above with Nos GAL4-VP16, which directs transcription in the germarium and subsequent stages of oogenesis. Females expressing Upf1-Nos produce essentially no eggs and bear rudimentary ovaries; in contrast, females expressing Upf1, Nos, or Upf1-Nos^L7^ appear to produce normal numbers of eggs. As explained in more detail below, we wished to harvest eggs at nuclear cycle 9 to take advantage of the prolonged transcriptional quiescence flanking the oocyte-egg transition (Fig 1A), and therefore asked whether use of other GAL4 drivers to direct expression of Upf1-Nos would permit egg development. We found this to be the case with a maternal *alpha*Tub> GAL4 driver, which directs expression throughout much of oogenesis and was used in all subsequent experiments.

Although females expressing Upf1-Nos lay eggs, virtually none hatch. In a small pilot study, 0/245 Upf1-Nos eggs hatched, whereas between 80 and 95% of Upf1, Nos, or Upf1-Nos Nos^L7^ control eggs hatched. Since expression of Upf1-Nos is exclusively maternal, we expected that defects might be evident early in embryonic development. Therefore we harvested 0-3 hour-old embryos and examined the distribution of nuclei by staining with DAPI to assess developmental progress. Unlike the three control genotypes, a large fraction of Upf1-Nos embryos have a single nucleus and are either not fertilized or fail to commence nuclear division cycles; we have not investigated this further, and restricted subsequent analysis to fertilized embryos between nuclear cycles 2-13. Almost all the Upf1, Nos, and Upf1-Nos Nos^L7^ embryos exhibit normal distributions of syncytial cleavage stage nuclei (94% - 100%, see S1 Data for details). In contrast, 63% of Upf1-Nos embryos have abnormal nuclear distributions, with irregularly spaced nuclei and variable DNA content, based on the intensity of DAPI fluorescence (e.g., Fig. 1C).

We next asked whether the Upf1-Nos embryos that fail to hatch exhibit interpretable developmental defects by examining their body plan, as revealed by pattern in the epidermal cuticle secreted late in embryonic development. 97% of Upf1-Nos eggs were “empty,” without detectable cuticle, probably because they die earlier in development (Fig. 1B). However, among the 3% of Upf1-Nos embryos that did secrete cuticle, 2 had normal segmentation and 14 revealed a striking bicaudal body plan, in which normal head and thoracic segments are replaced by a mirror image duplication of posterior abdominal segments (Fig. 1B). Bicaudal development is diagnostic for simultaneous inhibition of Bcd and Hb activity (33–35). None (0/862) of the Upf1, Nos, or Upf1-Nos Nos^L7^ control embryos exhibited segmental polarity reversals, much less a mirror image duplication of the abdomen. As shown in Fig. 1B, most (80%-95%) of the control embryos secreted cuticle that revealed body plans with normal antero-posterior polarity and, at most, minor defects; the remaining embryos failed to secrete cuticle.

Even though only a few Upf1-Nos exhibited bicaudal body plans, the phenotype is sufficiently characteristic to strongly suggest that Upf1-Nos inhibits activity of both Hb and Bcd. *bcd* is transcribed exclusively maternally, while *hb* is transcribed both maternally and later, zygotically (under the control of Bcd). For simplicity, we therefore focused on examining the effects of Upf1-Nos on *bcd* mRNA by in situ hybridization in early cleavage stage embryos. Most (82%) of the Upf1-Nos embryos that were either unfertilized or in the first nuclear division cycles have a normal level of *bcd* mRNA at the anterior (Fig. 1C); this was also the case for most (95%) control Upf1-Nos^L7^ embryos in cycles 0-2. But we observed a novel phenotype in Upf1-Nos embryos that we judged to be approximately in cycles 9-10 based on nuclear density: an absence of detectable *bcd* mRNA (Fig. 1C). In contrast, Upf1-Nos^L7^ embryos in cycles 9-10 have normal levels and distributions of *bcd* mRNA at the anterior pole.

In summary, the vast majority of control Upf1, Nos, and Upf1-Nos^L7^ embryos develop normally, which is consistent with subsequent analysis of their transcriptomes (below). In contrast, Upf1-Nos embryos are abnormal: many fail to either fertilize or commence nuclear division, subsequent nuclear divisions are abnormal, only a handful escape the deleterious early effects of Upf1-Nos and progress through to late development, and none hatch. Most importantly, two phenotypes--the absence of *bcd* mRNA during early development and the bicaudal body plan of Upf1-Nos embryos--support our premise that the chimeric protein would degrade Nos-regulated mRNAs.

### Upf1-Nos depletes 39% of maternal mRNAs

To identify targets of Upf1-Nos in the entire transcriptome, we prepared triplicate samples from 4 experimental embryos (Upf1, Nos, Upf1-Nos, Upf1-Nos^L7^) as well as wild type embryos as a control. Embryo collections were soft-fixed in methanol, stained with DAPI, and manually sorted in the fluorescence microscope to identify embryos at nuclear cycle 9-10, or in the case of Upf1-Nos, embryos that appeared to be approximately the same age despite an abnormal distribution of nuclei (as in Fig. 1C). Because transcription in the germ line ceases at stage 10B of oogenesis and resumes only during the MZT (maternal zygotic transition) beginning at nuclear cycle 10, pre-existing Upf1-Nos (or the other ectopically-expressed regulators) in the collected embryos have had the opportunity to act on the maternal transcriptome for ∼9-10 hours without synthesis of new mRNA. Prior to the transcriptional “shutoff,” Upf1-Nos might cause changes to the transcriptome via both direct and indirect mechanisms; we hoped that direct action of Upf1-Nos during the period of transcriptional quiescence would have a predominant effect on transcriptome composition in embryos harvested at nuclear cycle 9. RNA was prepared from all the genotypes, prepared for RNAseq, and analyzed by standard methods.

We first assessed the quality of our samples. The absence of detectable mRNAs encoded by genes that comprise the first zygotic response to maternal patterning gradients (e.g., *Kr*, *gt*, *kni*) supports the idea that embryos were in fact collected prior to the MZT (see S2 Data). Reducing the complexity of all 15 samples by Multi-dimensional Scaling reveals that biological replicates are similar to each other and that the control samples are also similar to each other (S1 Fig). In contrast, the Upf1-Nos samples are markedly different from all three UAS controls and wild type, consistent with their unusual embryonic phenotypes (Fig. 1).

We next compared the transcriptomes of each experimental genotype to wild type to identify genes with altered expression, arbitrarily setting a minimum two-fold threshold change and a FDR < 0.05. For simplicity, we describe data in which all mRNA isoforms for a gene are binned; analysis of individual mRNA isoforms yields broadly similar results (S2 Fig). We initially found that a small fraction of the maternal transcriptome is depleted (2.4%) or enriched (2.1%) upon expression of Upf1 alone (Fig 2A). Because we were interested primarily in mRNAs that are targeted by the Nos portion of the Upf1-Nos chimera, we excluded Upf1-regulated genes from subsequent analysis of Upf1-Nos and Upf1-Nos^L7^.

**Fig 2.**
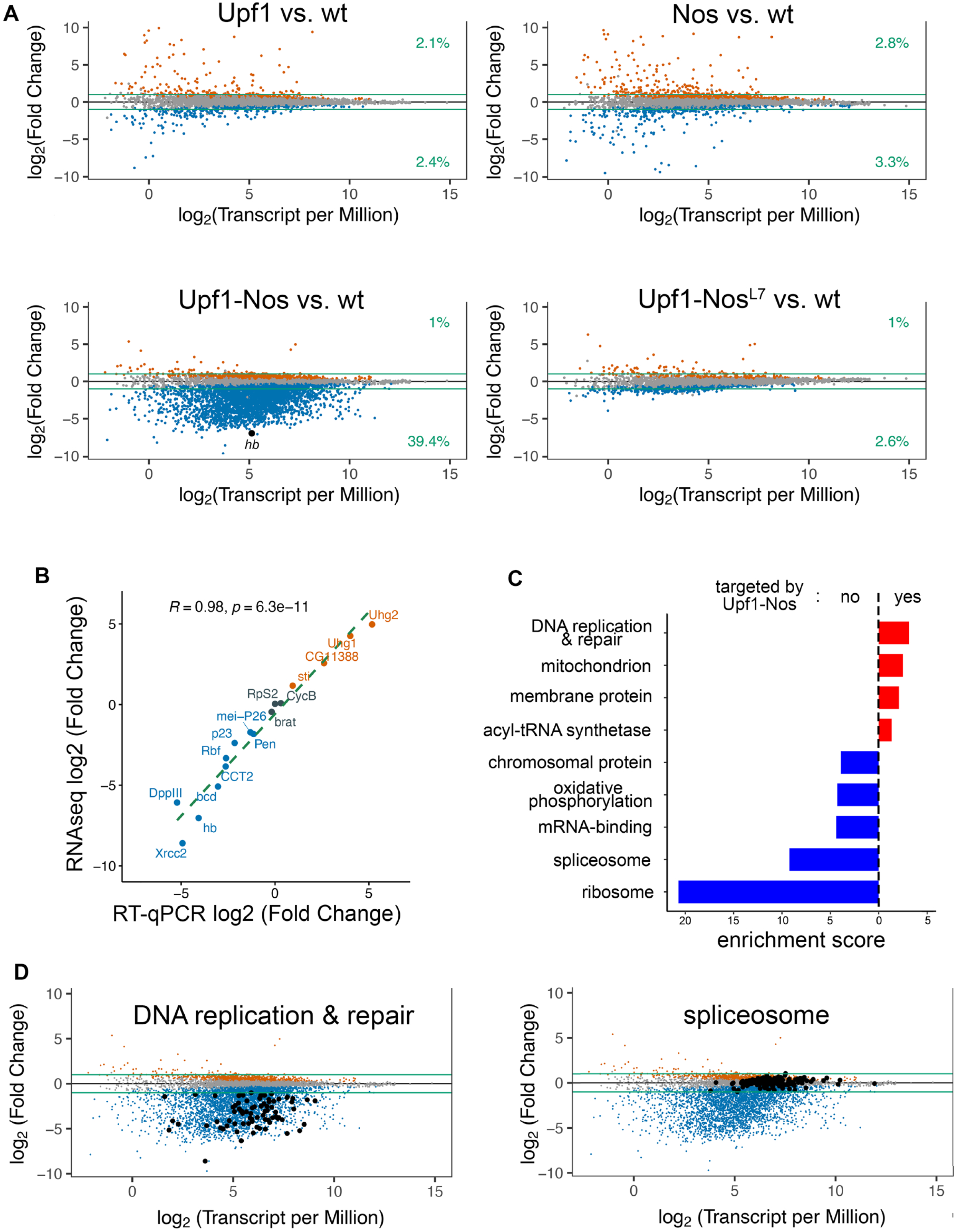
Upf1-Nos targets a large fraction of the maternal transcriptome. A. For each of the mis-expressed proteins, the plot shows the difference in mRNA abundance in comparison to wt (log_2_ Fold Change) as a function of the average mRNA abundance (log_2_ Transcripts per Million reads) for embryonic samples collected at nc 9-10. Each dot represents a binned collection of mRNA isoforms for a single gene; analysis of individual isoforms is in S2 Fig. Genes altered by expression of Upf1 are excluded from the analysis of Upf1-Nos and Upf1-Nos^L7^ data. Gene-binned mRNAs significantly depleted and enriched are indicated in blue and orange, respectively. The green lines mark log_2_ ± 1, and the figures at the right indicate the fraction of the (gene-binned) transcriptome that is significantly changed and above or below the arbitrary twofold cutoff. *hb* is highlighted in the Upf1-Nos plot. B. Correlation between the difference in mRNA abundance (Upf1-Nos vs. wt) measured by RNAseq and by RT-qPCR for a collection of genes, which span an ∼1000-fold range in expression level in wt. C. The results of DAVID analysis reveal gene sets enriched among Upf1-Nos-targeted mRNAs (red) and non-targeted mRNAs (blue), which are plotted vs. their enrichment scores (ES). Shown are annotation clusters with highest ES for which at least one Category has a Benjamini adjusted *p*-value < 0.05. For clarity, three clusters that satisfy these criteria and are enriched among non-targeted mRNAs with ES between 2.3 and 1.9 were omitted (see S2 Data for details). D. Examples of enriched gene sets, highlighted on the Upf1-Nos vs. wt plot from A.

As shown in Fig 2A, the expression of Upf1, Nos, or Upf1-Nos^L7^ has relatively little effect on the transcriptome. The proportion of significantly up- and down-regulated genes is approximately the same for Upf1 and Nos, and the proportion of the transcriptome depleted by Upf1-Nos^L7^ is only modestly greater than the proportion of up-regulated genes. In striking contrast, >39% (n = 2596) of the genes encoding maternal mRNAs are depleted more than twofold by Upf1-Nos. We note that the large, asymmetric effect of Upf1-Nos on the transcriptome results in over-sampling of non-targeted mRNAs; as a result we normalized the data using an average derived from 20 highly expressed mRNAs (most of which encode ribosomal proteins), as described in Methods and S2 Data. To test the normalization and obtain an independent measure of changes in abundance due to expression of Upf1-Nos, we measured the levels of 16 mRNAs by RT-qPCR; these were chosen to span a log_2_ = 15-fold change in abundance. As shown in Fig. 2B, the two methods show excellent agreement, with a Pearson’s correlation R value of 0.98. The absence of a significant population of enriched mRNAs in the Upf1-Nos data supports a premise of our experiment—that tethering Upf1 to mRNAs via Nos during the transcriptionally quiescent period preceding sample collection would lead to their degradation via NMD and that this degradation would predominate in shaping the composition of the transcriptome. We assume that indirect effects (i.e., up-regulation of some mRNAs as a secondary consequence of the action of Upf1-Nos on other, directly-targeted mRNAs) are minimized in the absence of concurrent transcription.

Several observations support the idea that, although Upf1-Nos targets a large portion of the transcriptome, it does so specifically. First, the inactivity of Upf1-Nos^L7^ argues that the wild type chimera acts in collaboration with Pum to specifically target NRE-bearing mRNAs, a point we return to in the following section. Second, three of the four mRNAs previously shown to be bound and regulated by Nos--*hb*, *bcd*, and *alpha-Importin* (36,37)-- are targeted by Upf1-Nos; moreover, *hb*, which is the critical Nos target in the early embryo, is one of the most highly depleted mRNAs (highlighted in Fig 2A). The exception is *CycB* mRNA, which is bound by Nos in collaboration with Pum and repressed in the PGCs (1,7), but not targeted by Upf1-Nos. [We note that, for reasons that are not clear, Nos is necessary but not sufficient to repress most of the *CycB* mRNA in these experiments which derives from the prospective somatic cytoplasm (1)]. Third, if Upf1-Nos acted promiscuously, due to high non-specific binding (for example), it would preferentially target mRNAs with longer 3’-UTRs; in fact mRNAs targeted by Upf1-Nos have on average slightly shorter 3’-UTRs than non-targeted mRNAs (S3 Fig).

The depletion of such a large fraction of the transcriptome by Upf1-Nos suggests that Nos may regulate thousands of mRNAs in Drosophila, in contrast to the hundreds regulated by Nos in C. elegans (10). Since such a large fraction of the transcriptome is targeted by Upf1-Nos, it is unclear whether it is meaningful to identify preferentially targeted pathways. Nevertheless, clustering analysis of gene function via DAVID (38,39) reveals that a number of pathways are significantly enriched among the pool of genes targeted by Upf1-Nos as well as the pool of untargeted genes (Fig 2C). Among Upf1-Nos targeted genes, only four clusters have gene subsets with significant Benjamini-adjusted *p*-values; the most highly enriched of these subsets encodes factors involved in DNA replication and repair (Enrichment Score [ES] = 3.1, Fig 2C), perhaps accounting for the aberrant replication seen in Upf1-Nos embryos (Fig. 1C). An independent search for Drosophila genes associated with the Gene Ontology terms “DNA replication” and “DNA repair” revealed that 79/79 are depleted, as shown in Fig 2D. Among genes <u>not</u> targeted by Upf1-Nos, constituents of the ribosome (ES= 20.7) and spliceosome (ES = 9.2) are even more significantly enriched (Figs 2C,D). We conclude that while Upf1-Nos selectively targets a few biological processes/pathways with modest preference, it acts globally to deplete maternal mRNAs that encode factors distributed across most biological functions.

### Upf1-Nos acts primarily in conjunction with Pum

Three lines of evidence support the idea that, in targeting a large fraction of the maternal transcriptome, Upf1-Nos acts in collaboration with Pum, much as native Nos and Pum collaboratively repress *hb* mRNA.

First, Upf1-Nos^L7^ is almost inert, changing the transcriptome to the essentially same minor extent as does Upf1 (Fig 2A). The C-terminal tail of Nos that is altered in the L7 mutant is specifically required to promote cooperative binding of Pum and Nos to the NREs in *hb* mRNA.

Second, a set of 538 ovarian mRNAs that co-purify with affinity-tagged Pum (27) is enriched among mRNAs depleted by Upf1-Nos [*p*-value = 0.03 by the CAMERA competitive test of (40), Fig 3A].

**Figure 3.**
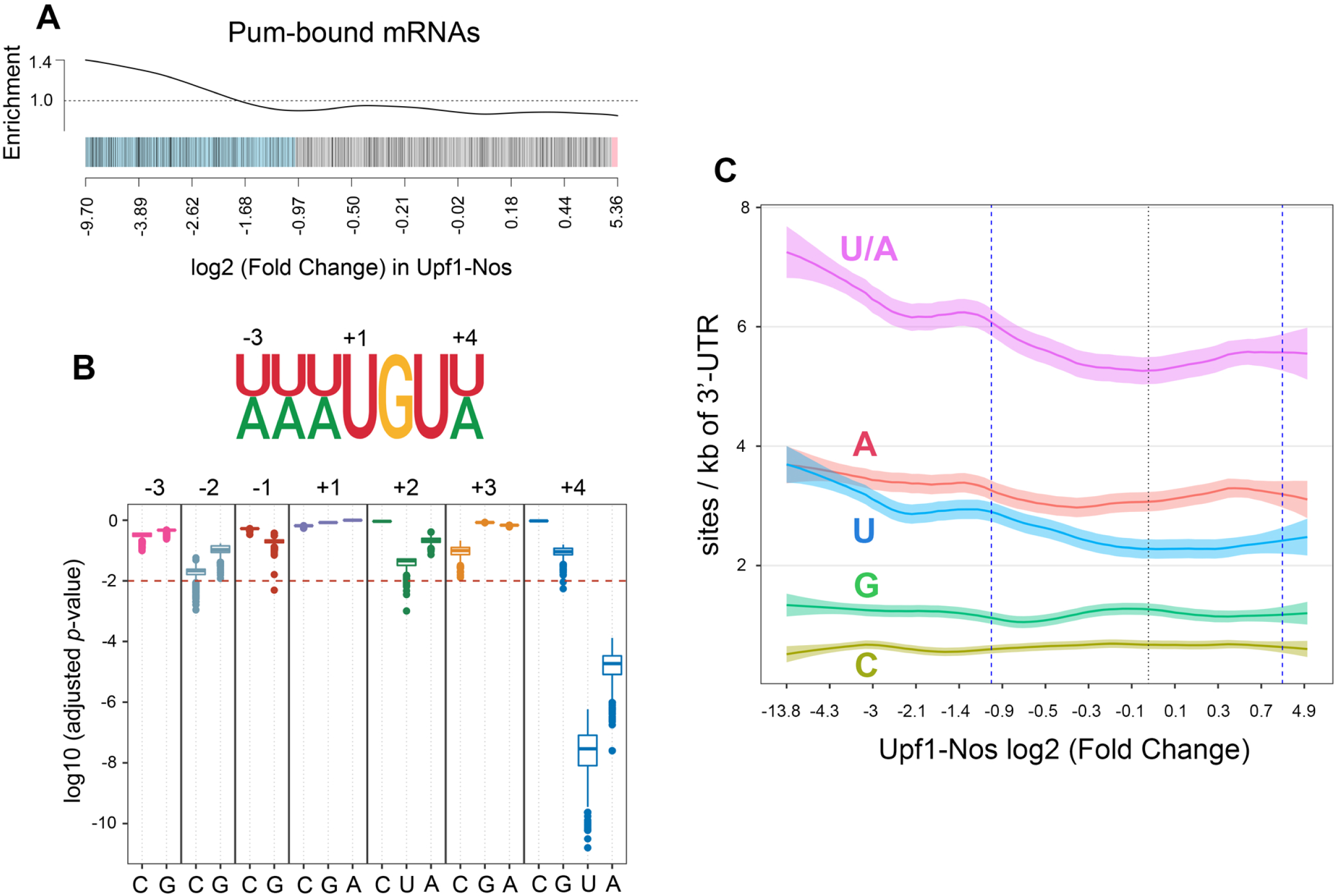
Pumilio mediates most of the activity of Upf1-Nos. A. Barcode plot showing enrichment of mRNAs depleted by Upf1-Nos among the mRNAs bound by transgene-encoded Pum RNA-binding domain in ovarian extracts identified by Gerber et al. (2006). Each mRNA is represented by a line; mRNAs depleted and enriched > log_2_=1 are highlighted in blue and red, respectively. Here and in subsequent barcode analyses, the x-axis is non-linear. B. A modified NRE motif (shown above) is enriched among the 3’-UTR sequences of mRNAs depleted by Upf1-Nos. The plot below is of adjusted *p*-values (Hommel) for the significance of the enrichment on the y-axis of each motif variant displayed along the x-axis. Enrichment was calculated as described in the text, and each boxplot displays the distribution of *p*-values from 1000 trial comparisons between the fraction of 3’-UTRs bearing a motif in targeted vs. non-targeted mRNAs. C. Local regression analysis displays the density of NRE variants at position +4 in 3’-UTR sequence plotted against the log_2_ (Fold Change) in Upf1-Nos vs. wt (e.g., Fig 2A). Here and in subsequent local regression analyses, the x-axis is non-linear, light shading marks 95% confidence windows, the light dashed line marks log_2_ = 0, and the heavier dashed lines mark log_2_ ± 1. Tests of whether the various NRE derivatives are more abundant among targeted 3’-UTRs (one-sided Wilcoxon) yield the following *p*-values: (U or A) < 2.2e-16, (A) 5.4e-07, (U) < 2.2e-16, (G) 0.057, (C) 1.

Third, the 3’-UTRs of mRNAs targeted by Upf1-Nos are enriched for a modified Nos+Pum binding motif, 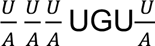, in which position +4 is either U or A. Several observations support this conclusion. We first asked whether the canonical Nos+Pum motif 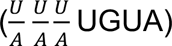 is enriched among 3’-UTRs targeted by Upf1-Nos, comparing the fraction of 3’-UTRs bearing the motif between targeted mRNAs and randomly selected subsets of non-targeted mRNA (chosen to have the same number of sequences and a similar distribution of 3’-UTR lengths). As shown at the far right of Fig 3B, the fraction of targeted 3’-UTRs bearing a canonical site is significantly greater than the fraction of non-targeted UTRs (median *p*-value of 1000 trials = 1.9e-05). As controls we asked whether singly-substituted mutant derivatives at each of the seven motif positions are also enriched. All of the derivatives but one appear to be “mutant” sites not significantly enriched in targeted 3’-UTRs; the exception is 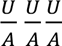 UGUU which, in fact, is enriched to a greater extent than the canonical site (Fig 3B). We next determined that the average number of modified Nos+Pum motifs is greater among 3’-UTRs targeted by Upf1-Nos than among non-targeted 3’-UTRs (*p*-value < 12.2e-16 by Wilcoxon rank sum test). Finally, localized regression analysis graphically reveals a correlation between the density of 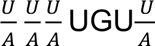 sites in the 3’-UTR and the extent of depletion by Upf1-Nos (Fig 3C). All of these observations are consistent with work described elsewhere showing that Nos+Pum bind efficiently to NREs with either U or A at position +4 (41).

Taken together, the evidence described above supports the idea that Upf1-Nos is primarily recruited to its targets by Pum in a manner similar to their joint recruitment to the *hb* NRE. We were unable to test Pum dependence directly, as *pum* mutant flies that also express Upf1-Nos are subviable, and females produce very few eggs.

### Upf1-Nos may be recruited to a minor fraction of its targets by Bruno

Although Upf1-Nos^L7^ is largely inactive (as described above), a small cohort of 178 genes is depleted > twofold upon expression of this mutant chimera. Localized regression analysis reveals no significant correlation between the density of NREs and the extent of depletion (Fig 4A, *p*-value = 0.087 by Wilcoxon test), and thus depletion is unlikely to be the result of inefficient recruitment of the mutant Nos moeity to NREs via Pum. We therefore considered the possibility that another RNA-binding protein might recruit Upf1-Nos ^L7^ to this minor fraction of the transcriptome.

**Figure 4.**
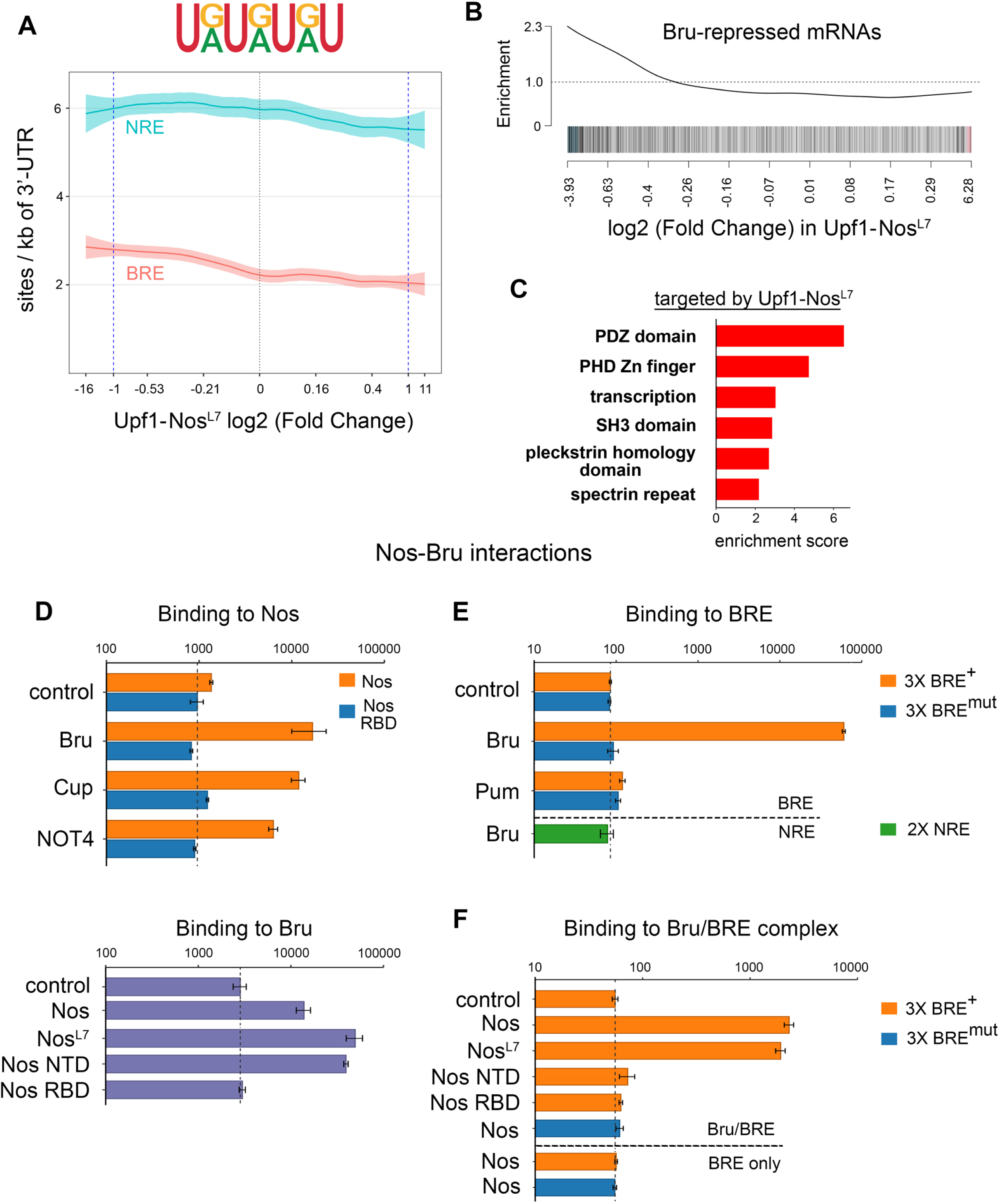
Upf1-Nos targets a small portion of the transcriptome via Bruno. A. Similar to Fig 3C, local regression analysis displays the density of BRE and NRE sites in 3’-UTR sequence plotted against the log_2_ (Fold Change) in Upf1-Nos^L7^ vs. wt (e.g., Fig 2A). Statistics for enrichment among mRNAs depleted > 2-fold reported in the text. B. Barcode enrichment plot displaying mRNAs de-repressed upon inhibition of Bru expression in the ovary (42) plotted vs. the log_2_ (Fold Change) in Upf1-Nos^L7^ vs. wt (e.g., Fig 2A). C. DAVID analysis reveals enrichment of gene sets among mRNAs targeted by Upf1-Nos^L7^. D-F. Yeast interaction assays show that both Nos and Nos^L7^ can be recruited to RNA-bound Bru. In each panel, the binding of AD-fusions to various factors (y-axis) to a bait (title above each bargraph) is measured in arbitrary light units of β-galactosidase (x-axis, log_10_ plot). For the “control” entry in each bargraph, yeast were transformed with plasmids that express the bait indicated in the title and an empty vector encoding AD only; the background level of β-galactosidase is also indicated with a dashed line. D. Two-hybrid experiments showing protein-protein interactions between Nos and Bru (above) and Bru and Nos (below). We are unable to assess interaction with DBD fused to the Nos NTD, as the resulting fusion “auto-activates” transcription of the reporter without an AD-fusion partner. E. Three-hybrid experiments showing specific binding of Bru to RNA bearing wt BREs. Below the horizontal dashed line is an additional control showing no significant binding of Bru to NREs. F. Four-hybrid experiments showing binding of Nos and Nos^L7^ to Bru-bound RNA. Below the horizontal dashed line are additional controls showing no binding of Nos in the absence of Bru to RNA bearing either wt or mutant BREs. Note that the background level of β-galactosidase is higher in the yeast strain used for 2-hybrid experiments than in the strain used for 3- and 4-hybrid experiments.

To identify such a putative novel Nos co-factor, we used the DREME tool (43) to search for motifs overrepresented among 3’-UTR sequences of mRNAs depleted by Upf1-Nos^L7^. The search revealed enrichment of the motif 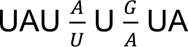, which is similar to one of the binding sites for Bruno: the Bruno Response Element (BRE) 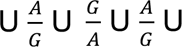 (44). Localized regression analysis further shows that mRNAs depleted ≥ twofold by Upf1-Nos^L7^ are enriched in BREs (Fig 4A, *p*-value < 2.2e-16 by Wilcoxon rank sum test).

Bruno (Bru) is a translational repressor that binds a number of RNA motifs via its three RNA recognition motifs (RRMs) (45). mRNA regulatory targets of Bru in the ovary have been identified by virtue of increased polysome association following RNAi-mediated inhibition of Bru expression (42). If Bruno mediates the depletion of mRNAs by Upf1-Nos^L7^ in our experiments, we would expect significant overlap with the Bru-regulated mRNAs identified by Rangan and colleagues. This is indeed the case: mRNAs depleted by Upf1-Nos^L7^ are enriched for Bru-regulated targets (*p*-value = 7.2e-6, CAMERA competitive test), as revealed in a barcode enrichment plot (Fig. 4B). Functional clustering of gene analysis reveals a number of enriched gene sets among Upf1-Nos^L7^ targets; these encode membrane cytoskeleton factors or regulators (PDZ, SH3, pleckstrin domain proteins, spectrin repeat proteins), at least one of which (β-spectrin) plays a key role in GSCs and differentiation of their progeny (Fig 4C and S3 Data).

We next asked whether Bru can recruit Nos and Nos^L7^ to RNA. Previous experiments have shown that Nos and Bru proteins interact (46,47). To further characterize the Bru-Nos interaction, we performed yeast two-hybrid experiments, the results of which are shown in Fig 4D. We find that a fragment of Bru interacts with full-length Nos, stimulating LacZ reporter activity to a slightly greater extent than does interaction with either of two known Nos effectors, Cup and NOT4 (Fig 4D above) (1,48). In reciprocal experiments with DNA-binding domain (DBD) and transcriptional activation domain (AD) fusions swapped, we observe binding of both Nos and Nos^L7^ to Bru, demonstrating that the Bru-Nos interaction does not depend on an intact Nos C-terminal tail. Consistent with this observation, Bru interacts with the Nos N-terminal domain (NTD) but not its C-terminal RBD (Fig 4D below).

To directly test whether RNA-bound Bru can recruit Nos to RNA, we next used yeast three-hybrid RNA-binding experiments to show that Bru binds specifically to a synthetic RNA sequence with 3 copies of the canonical BRE, but not to RNAs with either 3 copies of a mutant BRE or 2 copies of a NRE (Fig 4E). Then in yeast four-hybrid experiments similar to those used to show Pum-dependent recruitment of Nos to NREs (23), we find that RNA-bound Bru can recruit either Nos or Nos^L7^ to the RNA (Fig 4F). Recruitment is dependent on both the NTD and RBD of Nos, raising the possibility that protein-protein interactions between Bru and Nos as well as protein-RNA interactions between Nos and the RNA are required for formation of the ternary Bru/Nos/RNA complex.

Although we initially identified Bru as a mediator of the activity of Upf1-Nos^L7^, the interaction experiments in Fig 4 (D-F) show that Bru does not interact specifically with the L7 mutant form of the protein but also to a similar extent with wild type Nos. It therefore seems likely that Bru mediates targeting of the same small subset of mRNAs by both Upf1-Nos^L7^ and Upf1-Nos, but that Pum-mediated targeting of the much larger set of mRNAs by the latter obscures the contribution by Bru. Consistent with this idea, neither BREs (*p*-value = 0.129 by Wilcoxon rank sum test) nor Bru-repressed mRNAs (*p*-value = 0.084 by CAMERA competitive test, S4 Fig) are enriched among mRNAs targeted by Upf1-Nos.

In summary, we have shown that Bru can recruit Nos or Nos^L7^ to specific RNA sequences, supporting the idea that Bru mediates targeting of a minor subset of the maternal transcriptome by Upf1-Nos^L7^ (and presumably contributing to depletion of the same minor subset by Upf1-Nos). The mechanism by which the Nos-Bru-BRE complex forms is independent of the integrity of the Nos C-terminal tail and thus different from cooperative binding of Nos+Pum to the *hb* NRE.

### mRNAs targeted by Upf1-Nos in transgenic animals are repressed in wildtype ovaries and embryos by Nos

We next asked whether mRNAs depleted by Upf1-Nos might be regulated by endogenous Nos in wildtype animals. As a first step to addressing this question, we re-analyzed two studies of maternal mRNAs in late oogenesis and early embryogenesis, partitioning these into mRNAs either targeted or not targeted by Upf1-Nos.

In the first study, Lipshitz, Smibert, and colleagues (49) identified maternal mRNAs that are degraded in unfertilized eggs; by definition, the regulation they observed occurs in the absence of zygotic gene expression. Tadros et al. (2007) identified 1069 mRNAs whose abundance drops at least 1.5-fold as unfertilized eggs age. This set of destabilized maternal mRNAs is significantly enriched among our Upf1-Nos targets (*p*-value = 1.6e-11; Fig 5A). The embryos used in their experiments were unfertilized but otherwise wildtype, and therefore destabilization must be mediated by endogenous maternal factors.

**Figure 5.**
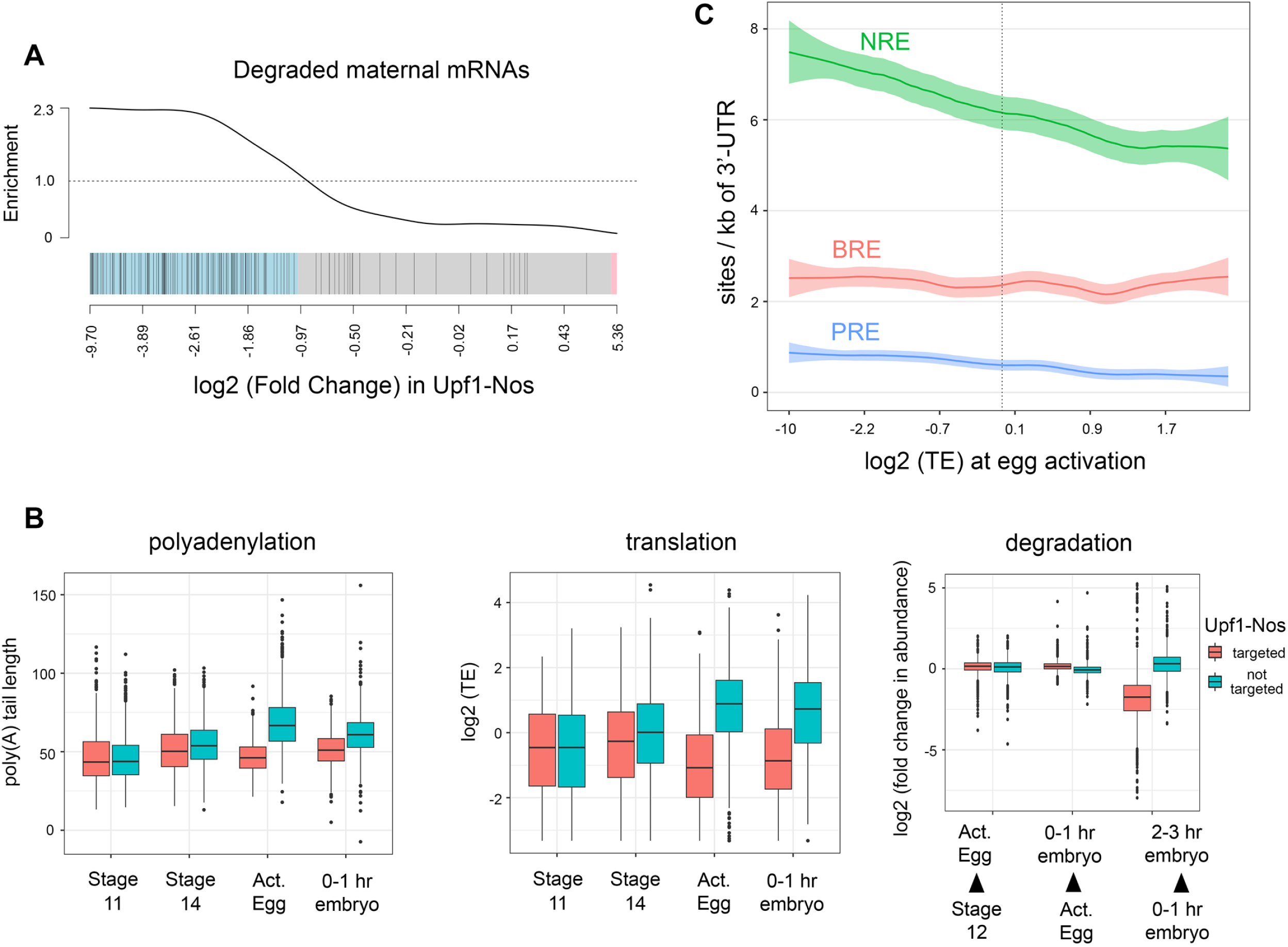
Maternal mRNAs targeted by Upf1-Nos are translationally repressed and degraded in wild type eggs. A. Barcode plot displaying enrichment of maternal mRNAs degraded in unfertilized eggs (49) vs. log_2_ (Fold Change) in Upf1-Nos vs. wt (e.g., Fig 2A). B. Maternal mRNAs expressed at nc 9-10 in the RNAseq experiments of Fig 2 were binned into those targeted (red) and not targeted (teal) by Upf1-Nos. Then, the poly(A) tail length and relative translational efficiency of each mRNA was extracted from the data of Eichorn et al. (2016) at the four stages of development indicated below: stage 11 and 14 oocytes, activated eggs, 0-1 hr embryos (before the onset of zygotic transcription). The plot at the right is of the change in abundance for each maternal mRNA between each of the three developmental transitions indicated below, again using the data of Eichorn et al. (2016). Statistical analysis in S4 Data. Note that zygotic transcription contributes significantly to mRNA abundance in the 2-3 hr embryo pool, but all other timepoints analyze purely maternal mRNAs. C. Local regression analysis of binding motifs in mRNA 3’-UTRs as a function of Translational Efficiency (TE) at egg activation, from the data of Eichorn et al. (2016). For this analysis, we chose the most highly expressed maternal isoform for each gene.

In the second study, Bartel, Orr-Weaver, and colleagues (50) characterized mRNAs in staged samples from wildtype ovarian egg chambers and timed embryo collections up to 6 hours after egg deposition, measuring abundance, poly(A) tail length, and translational efficiency (TE). Their work reveals comprehensive changes to the maternal transcriptome during an extended span of development that includes the oocyte-egg transition (e.g., the vertical dashed line in Fig 1A) and the maternal-zygotic transition (MZT), a period from approximately 1.5-3.0 hours of embryonic development during which many maternal mRNAs are degraded and zygotic transcription is activated. (For reference, the MZT begins shortly after nuclear cycle 10 in Fig 1A.) Analyzing their data, we find that mRNAs targeted by Upf1-Nos are regulated differently than are non-targeted mRNAs. Non-targeted maternal mRNAs undergo lengthening of their poly(A) tails at egg activation and concomitant enhanced translation, consistent with the observation that poly(A) tail length and TE are coupled until the onset of gastrulation (which occurs at 3 hours of embryonic development) (50). In contrast, Upf1-Nos targets undergo neither poly(A) tail lengthening nor translational enhancement (Fig 5B). The repressed Upf1-Nos targets are stable until later in the MZT (2-3 hours of development), when they are preferentially degraded (Fig 5B). The data likely underestimate the extent of degradation during the MZT, since some maternally transcribed mRNAs are re-transcribed zygotically, as discussed below.

Taken together, the data in Fig 5 suggest that, late in oogenesis, the large set of ∼2600 mRNAs targeted by Upf1-Nos (in transgenic animals) is translationally repressed in wild type animals during late oogenesis and degraded 2-3 hours later, during the MZT. We suggest that these regulatory events are a delayed response to the burst of Nos expression at stage 10B of oogenesis (Fig 1A), which represses targeted mRNAs and primes them for subsequent degradation in the embryo.

As a first test of the idea that Nos is responsible for repressing a large cohort of maternal mRNAs, we asked whether the modified NRE motif that mediates Nos+Pum binding is over-represented among maternal mRNAs that are translated inefficiently at egg activation. As shown in Fig 5C, localized regression analysis reveals a correlation between the translational efficiency and the density of modified NRE motifs in the 3’-UTR across the maternal transcriptome. In comparison with the 3’-UTRs of mRNAs that are not repressed, the modified NRE is enriched in the 3’-UTRs of repressed mRNAs (defined as TE < 1, *p*-value = 1.1e-8 by two-sided Wilcoxon test). As controls, canonical Pum binding sites and BREs are both present at lower density (Fig 5C) and are less enriched in repressed mRNAs (*p*-values of 7.3e-6 and 0.22, respectively).

All the work described above is based on the ectopic expression of Upf1-Nos, which behaves like a hyperactive version of Nos that efficiently degrades bound mRNAs. Part of the rationale for using Upf1-Nos is that, in the Drosophila embryo, Nos acts primarily to block translation and only secondarily to promote the degradation of targeted mRNAs (35,36,51,52). In C. elegans, RNAseq analysis of *nos* mutant PGCs successfully identified a cohort of maternal mRNAs that are destabilized by Nos in wt PGCs (10). This observation led us to ask (1) if we could detect Nos-dependent destabilization of maternal mRNAs, and (2) whether the same mRNAs are targeted by Upf1-Nos and by Nos. To address these issues, we collected embryos from wild type and *nos*^-^ females at 0-1 and 2-3 hours of development (i.e., before and after the onset of zygotic transcription, respectively), and measured the abundance of maternal mRNAs via RNAseq. In analyzing the data, we ignored mRNAs that are transcribed zygotically but not maternally, since these are not exposed to and thus cannot be regulated by Nos.

As shown in Fig 6A, at 0-1 hrs, the wt and *nos*^-^ mRNA populations are similar in composition, with only 3.5% of the maternal transcriptome significantly enriched >log_2_=0.5-fold in *nos*^-^ embryos. A somewhat greater fraction of the maternal transcriptome (5.6%) is depleted in *nos*^-^ embryos at this stage of development, presumably due to indirect effects of Nos on gene expression earlier, during ovarian development (since Nos is thought to act primarily as a repressor). However, after the onset of the MZT, 18.5% of the maternal transcriptome is stabilized or up-regulated in 2-3 hr *nos*^-^ embryos (Fig 6A). A rank-ordered barcode display of mRNAs stabilized in the absence of Nos reveals a correlation with mRNAs targeted by Upf1-Nos (*p*-value = 3.8e-14, Fig 6B left). In fact, 86% (1019/1185) of the mRNAs stabilized in the absence of Nos are depleted by Upf1-Nos (Fig 6B middle). Taken together, the complementary loss- and gain-of-function results of Figures 6 and 2 indicate that Nos regulates at least these 1019 common mRNA targets.

**Figure 6.**
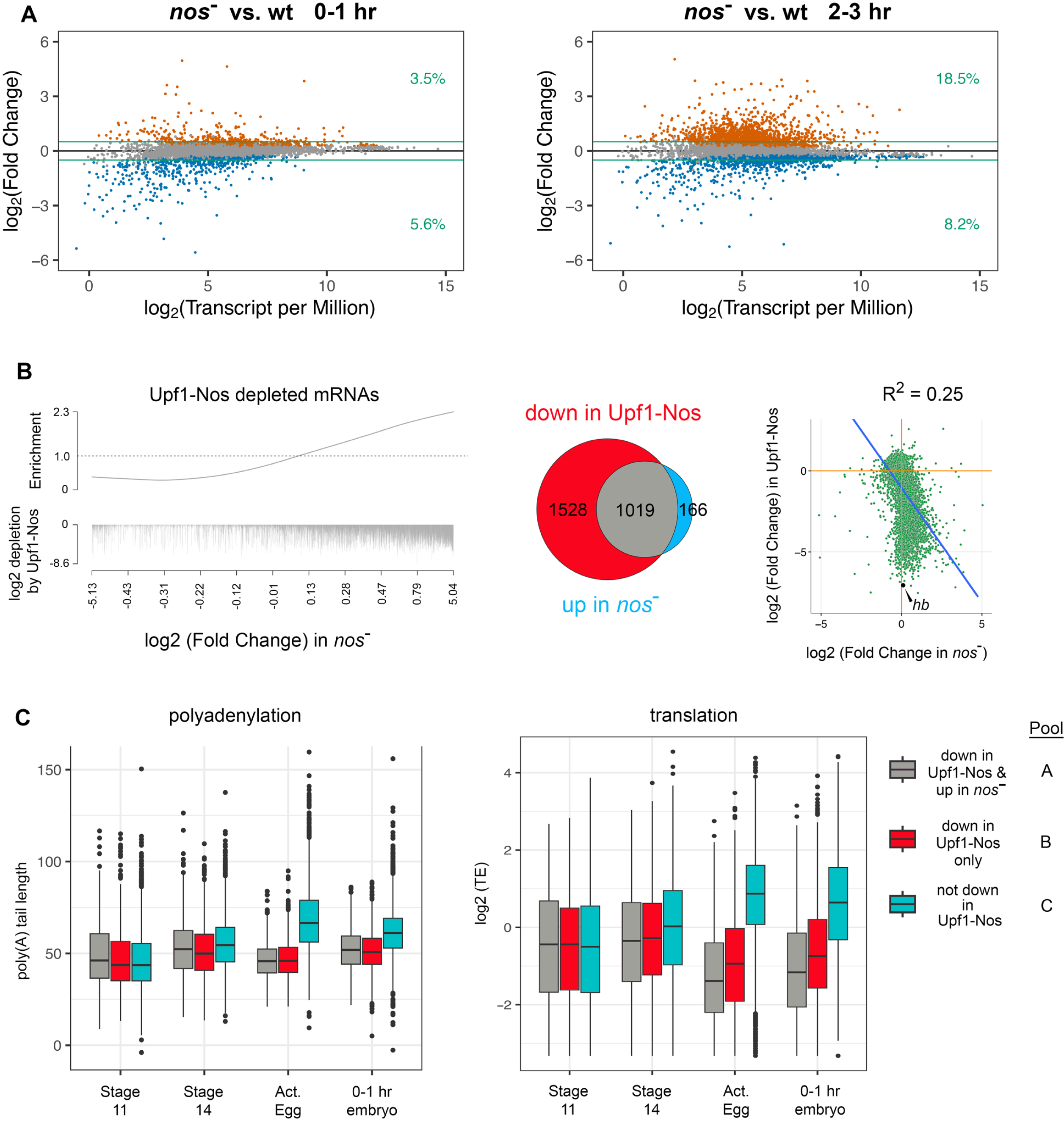
Regulation of maternal mRNAs by endogenous Nos in wild type oocytes and embryos. A. Results of RNAseq analysis, comparing the abundance of maternal mRNAs between wild type and *nos*^-^ embryos at 0-1 and 2-3 hr of embryonic development. Only maternal mRNAs (based on the manually sorted embryos used in the RNAseq experiments of Fig 2) are shown. The green lines mark log_2_ ± 0.5, and the figures at the right indicate the fraction of the (gene-binned) transcriptome that is significantly changed and above or below the arbitrary cutoff, which is smaller than the 2-fold cutoff used in Fig 2 since the magnitude of destabilization by Nos is generally smaller than the magnitude of degradation by Upf1-Nos. mRNAs stabilized or destabilized in the absence of Nos at an FDR < 0.05 are orange and blue, respectively. B. The barcode (left) shows mRNAs depleted by Upf1-Nos (Fig 2) plotted vs. the log_2_ (Fold Change) in *nos*^-^ vs. wt at 2-3 hr (e.g., Fig 6A). Each depleted mRNA is a hashmark, the length of which indicates the extent of depletion by Upf1-Nos; this plot shows a poor correlation between the extent of degradation by Upf1-Nos and by Nos (in wt embryos; see also the scatterplot to the right). The Venn diagram (middle) shows that most of the maternal mRNAs stabilized in the absence of Nos are degraded by Upf1-Nos. The scatterplot (right) shows the correlation between the extent of degradation by Upf1-Nos and stabilization in the absence of Nos. The canonical Nos substrate *hb* mRNA is highlighted as an extreme example, in which zygotic re-expression between 2-3 hrs almost completely obscures Nos-mediated regulation of maternal *hb* mRNA. In all these comparisons we analyze only the 6399 mRNAs common to the two datasets (e.g., Figs 2A and 6A). C. Maternal mRNAs were divided into three pools: down in Upf1-Nos and up in *nos*^-^ at 2-3 hrs (Pool A = the 1019 mRNAs of the Venn diagram above); down in Upf1-Nos only (Pool B = the 1528 mRNAs of the Venn diagram), the remaining 3852 maternal mRNAs untargeted by Upf1-Nos (Pool C). These bins were used to parse the data of Eichorn et al. (2016), as described for Fig 5. Statistical analysis in S5 Data.

What then of the 1528 maternal mRNAs that are regulated by Upf1-Nos but apparently not by Nos (Fig 6B middle)? One possibility is that they are repressed but not degraded by Nos. Another possibility is that they are repressed and degraded by Nos, and then re-expressed zygotically during the 2-3 hour window of embryonic development. At this stage, Nos is confined to the PGCs (which constitute only a small fraction of the volume of the embryo), and therefore essentially no zygotically re-expressed mRNAs will suffer regulation by Nos even if they might be susceptible. This is not a confounding issue for our analysis of Upf1-Nos activity, since mRNA was harvested before the onset of transcription. However, zygotic re-expression likely does mask regulation of mRNA stability by Nos, thereby accounting, at least in part, for the modest correlation between the extent of regulation by Upf1-Nos and by Nos (R^2^ = 0.25, Fig 6B right). For example, the canonical Nos target *hb* (highlighted in the scatter plot in Fig. 6B), which is among the most sensitive Upf1-Nos targets, is re-transcribed in the 2-3 hr window such that the net level of *hb* mRNA at 2-3 hrs of development is not significantly different in the presence or absence of Nos.

To estimate the potential influence of zygotic re-expression on the 2-3 hr samples ± Nos, we analyzed data from Eisen and colleagues (53). Their measurements of RNA abundance in precisely staged single embryos allows us to determine the fraction of maternal mRNAs in our experiments that are re-expressed zygotically within three pools: mRNAs targeted by both Upf1-Nos and Nos (pool A; 105/1019 = 10%); mRNAs targeted by Upf1-Nos but not Nos (pool B; 655/1528 = 43%); mRNAs targeted by neither (pool C; 760/3852 = 20%). There is a striking enrichment of zygotically re-expressed mRNAs in pool B (*p*-values < 2.2e-16 vs. pools A or C, respectively, by Fisher’s exact test). Taken together, these observations support the idea that, for many of the genes in pool B, Nos-dependent degradation of the maternal mRNA component might be masked by new transcription at 2-3 hrs. of embryonic development (e.g., as for *hb*).

We reasoned that we might observe a reduction in poly(A) tail length and TE of the 1528 mRNAs in pool B at earlier stages of development, before the potentially confounding contribution of zygotic transcription. To test the idea, we returned to the resource described by Eichorn et al. (2016), examining the fate of maternal mRNAs at various stages from late oogenesis through the first hour of embryonic development. Data were parsed into the three pools described above: (A) the 1019 mRNAs targeted both by Upf1-Nos and Nos, (B) the 1528 mRNAs targeted only by Upf1-Nos, and (C) the remaining 3852 mRNAs not targeted by Upf1-Nos. As shown in Fig 6C, the two pools containing Upf1-Nos-targeted mRNAs (A and B) are regulated in a similar fashion, particularly at egg activation: mRNAs in both pools are translationally repressed and suffer shortening of their poly(A) tails (albeit, to slightly different extents, S5 Data). In contrast, the mRNAs not targeted by Upf1-Nos (pool C) are translationally activated and undergo lengthening of their poly(A) tails.

Overall, we conclude that, at or around the oocyte-egg transition, endogenous Nos likely regulates most of the maternal mRNAs targeted by Upf1-Nos.

## Discussion

The main conclusions from the work presented above are that Nos regulates thousands of mRNAs in the maternal transcriptome and likely does so in concert with Pum, binding jointly to NRE sequences much as they do to the canonical NREs in *hb* mRNA. The latter idea is supported primarily from the relative inactivity of Upf1-Nos^L7^, which bears a defective C-terminal tail, a critical element of the intricate set of protein-protein and protein-RNA interactions that support formation of the Nos/Pum/*hb* NRE ternary complex. In theory, the Nos C-terminal tail might mediate similar interactions with another RNA-binding partner too. Although we cannot rule out such an idea, the enrichment of NREs among mRNAs depleted by Upf1-Nos (Fig 3) or repressed at egg activation (presumably by endogenous Nos, Fig 5) support the idea that most of the Nos tail-dependent activity is mediated via Pum as a partner.

A number of factors drove us at the outset to adopting the unconventional approach of using Upf1-Nos to target the transcriptome. First, we failed to achieve meaningful enrichment of *hb* mRNA in pulldown experiments from embryonic extracts expressing epitope-tagged Nos. This disappointing observation was mitigated by the modest enrichment of the canonical Nos+Pum targets *hb* or *bcd* mRNA in the Pum pulldown experiments of others (27,28). In these experiments, *hb* and *bcd* mRNA were enriched only between 2- and 3.5-fold, even though Pum on its own is a high-affinity, high-specificity binding protein. Given the modest enrichment for known Pum-bound mRNA targets, the high non-specific binding activity of Nos, and our insignificant enrichment of *hb* in pilot experiments, we were not encouraged to believe that we would be able to purify other, less stable Nos complexes formed on suboptimal NREs, for example. We also considered using UV-crosslinking to stabilize Nos/mRNA complexes in extracts; but crosslinking of the purified Nos RBD is inefficient (23). In contrast, in vivo expression of Upf1-Nos allowed us to achieve high activity and specificity: *hb* and *bcd* are depleted 131- and 34-fold, respectively, by Upf1-Nos, whereas upon expression of Upf1-Nos^L7^ *hb* is depleted only 1.5-fold and the level of *bcd* is not significantly changed. The large dynamic range of our experiment allowed the identification of thousands of less-sensitive targets, and over 1000 of these are verified by virtue of stabilization in the absence of Nos (Fig 6). Nevertheless, a caveat to the present work is that we have not shown that mRNAs regulated by either Upf1-Nos or Nos in vivo are directly bound by Nos in vitro. Nor have we identified specific sites within regulated mRNAs that mediate Nos action.

Although we have not identified the sites within regulated mRNAs that mediate Nos (or Upf1-Nos) action, analysis of the population of targeted mRNAs has shown that Nos+Pum preferentially recognize mRNAs with both UGUU and UGUA motifs. We observe a correlation between the density of such motifs in the 3’-UTR and sensitivity to Upf1-Nos, but have not been able to identify similar correlations between the total number of motifs or their position within the 3’-UTR. At one level this is surprising, since a single site is sufficient to confer regulation either in flies or in cultured cells, and the degree of regulation correlates roughly with the number of sites (22,37). Therefore, one might expect that mRNAs with many sites would be more sensitive, as the probability of occupancy by a Nos+Pum repressor complex would be greater. This is clearly not the case; for example, of the ten mRNAs in the transcriptome with the largest number of 3’-UTR UGUA motifs (26), only one is targeted by Upf1-Nos. The correlation that we do observe (Fig 3C) is consistent with the idea that higher site density promotes cooperative interactions among nearby bound complexes. This model is supported by experiments described elsewhere showing that the N-terminal domain of Pum interacts with itself (Wharton et al., 2023). Finally, the contribution of other RNA-bound regulators or RNA structure is too complex to currently be taken into account when considering the entire transcriptome, as has been pointed out previously (26).

Given the large number of regulated mRNAs, it is not surprising that expression of Nos protein is tightly controlled. During oogenesis and embryogenesis, much of this control is exerted via the 3’-UTR of its mRNA to control the timing and location of protein accumulation (31,54,55); we therefore intentionally designed all of the transgenes used in this work to express mRNAs bearing the native *nos* 3’-UTR. Our results suggest an additional level of control--negative autoregulation. Nos mRNA is significantly depleted (6.7-fold) by Upf1-Nos, and most of this regulation appears to be Pum-mediated (e.g., only 1.3-fold depletion by Upf1-Nos^L7^). The idea of negative autoregulation is consistent with earlier work on the activity of Nos at the neuromuscular junction, which hinted at Pum-dependent auto-regulation via non-canonical sites in the *nos* 3’-UTR (16).

Based initially on analysis of the low residual activity of Upf1-Nos^L7^, we argue above that Bru may recruit Nos (or Nos^L7^) to regulate a minor portion of the maternal transcriptome. Such a model could explain an intriguing aspect of Nos biology in the female germline stem cell and its progeny cystoblast, which differentiates into the cyst of 16 cells that ultimately forms the germline component of each egg chamber. In females bearing near-complete loss-of function *nos* alleles (e.g., *nos*^RC^), GSCs are specified normally but fail to maintain stem cell status following division: both daughter cells differentiate and the ovary is rapidly denuded of its germline (56). However, GSC function is maintained in the *nos*^L7^ mutant, suggesting that at least some of the essential Nos function at this stage is tail-independent and therefore not mediated by Pum (24).

Our results are consistent with the idea that Bru mediates part of the activity of Nos in early oogenesis, and does so whether the Nos C-terminal tail is intact or not. Bru is required both early in the larval gonad for normal development of the GSCs in the larval ovary and of their cystoblast progeny in the adult ovary (57). The definitive cellular marker for the GSC and the cystoblast is the spectrosome, an unusual intracellular sphere containing cytoskeletal proteins usually associated with the cellular membrane--alpha- and beta spectrin, spectraplakin (which links actin and microtubule filaments), and an adducin (encoded by *hts*), which mediates interactions between spectrin and actin (58–60). We find that the mRNAs encoding these proteins and functionally related cytoskeletal factors are enriched among Upf1-Nos^L7^ targets (Fig 4C). Taken together, these observations lead us to speculate that at least part of the function of Nos in maintaining stem cell identity in the GSCs may rely on Bru-mediated regulation of mRNAs encoding the spectrosome and other cytoskeletal components.

Part of the motivation for identifying novel Nos regulatory targets was the hope of gaining insight into its evolutionarily conserved role in the GSCs, where it maintains stem cell identity by unknown pathways, and the PGCs, where it regulates proliferation, contributes to selectively blocking transcription of somatic genes, and promotes escape from apoptosis. The large number of Nos targets we have identified suggest it is unlikely that Nos controls a single mRNA to maintain the GSCs in a manner similar to the singular regulation of maternal *hb* mRNA that drives abdominal patterning in the embryo. Translation is globally repressed in human hematopoeitic stem cells (61); perhaps Nos-mediated global repression contributes to maintenance of a semi-dormant state that promotes GSC maintenance. Alternatively, targeting of particular pathways could regulate GSCs. For example, our finding that DNA replication gene mRNAs are preferentially targeted by Upf1-Nos might depress fork speed, which has recently been shown to promote a totipotent state in mouse embryonic stem cells (62). Upf1-Nos preferentially spares ribosomal protein mRNAs, perhaps so that, following GSC cell division, the cystoblast daughter (in which Nos drops to a low level) can immediately commence differentiation. In addition, it is intriguing that Upf1-Nos spares proteins that regulate mRNA biology (e.g., those encoding RRM proteins), given the importance of post-transcriptional regulation in the GSC in particular and the germline in general. Further work will be required to determine whether any of this regulation contributes to GSC (or PGC) biology.

We initially analyzed the effects of Upf1-Nos on the maternal transcriptome at nc 9-10 largely because the preceding period of transcriptional quiescence provided a convenient method of analyzing post-transcriptional effects on mRNA levels in the absence of new synthesis. However, our findings suggest an unanticipated role for the burst of Nos expression at stage 10B of oogenesis, just prior to the transcriptional shutoff. Turnover of part of the maternal transcriptome is an essential aspect of embryonic development in Drosophila. Degradation is controlled by a number of factors (including Smg, Brat, ME31B, and mir-309; reviewed by (63) in successive waves during syncitial embryonic development. Our finding that mRNAs targeted by Upf1-Nos and/or Nos are preferentially hypo-adenylated and translationally repressed at egg activation (Fig 6) is consistent with the idea that Nos contributes to this degradative regulatory network. We favor the idea that the high level burst of Nos expression at oogenesis stage 10B represses thousands of mRNAs, priming them for further subsequent repression in the embryo by other factors and, ultimately, degradation. In theory, the nascent posterior gradient of Nos, which initially forms at oogenesis stage, could contribute as well. But given the difference in Nos protein levels at stages 10B and 13-14 (31), we favor a model in which the stage 10B burst is responsible for most of the coordinated regulation of poly(A) tail length, translational efficiency, and stability that is observed. In this scenario, Nos is responsible for regulating mRNA fate on two very different scales in the early embryo--repressing a single mRNA (*hb*) to govern abdominal segmentation and thousands of mRNAs to help direct turnover of the maternal transcriptome.

## Materials and methods

### Drosophila strains and methods

Plasmids used in this study are described in S6 Data. Transgenic lines were constructed by microinjection of *w*^1118^ embryos by standard methods. The following strains were used: *w*^1118^ as the wild type reference in RNAseq experiments; the maternal alpha-tubulin GAL4-VP16 driver (stock # 7062 from the Bloomington Drosophila Stock Center [BDSC]); trans-heterozygotes of *nos*^BN^ and Df(3R)Exel6183 (stock # 7662, [BDSC]) for the RNAseq experiment of Fig. 6. We initially tested 3 or more transgenic lines with the GAL4-VP16 driver for each construct, and observed only minor phenotypic differences among the 3 lines. We screened for lines expressing Upf1-Nos and Upf1-Nos^L7^ at essentially the same level by harvesting total RNA from collections of 0-2 hr embryos and measuring the chimeric mRNA by RT-qPCR using the appropriate primer pair in S2 Data. Fly stocks were maintained by standard methods and grown at 25°C. Embryos were collected on apple-juice agar plates smeared with yeast paste.

Early embryos for assessment of nuclear morphology (Fig 1) were prepared by dechorionating in bleach, harvesting by filtration, fixing in 4% formaldehyde under heptane for 20-30 minutes, removal of the aqueous phase and shocking into methanol. Embryos were then rehydrated into phosphate-buffered saline (PBS) containing 0.1% Tween-20 (PBT), and incubated with 5 µg/ml 4′,6-diamidino-2-phenylindole (DAPI) for 10 minutes, washed repeatedly with PBT, mounted in PBST containing 25% glycerol, and examined by epifluorescence using a Zeiss Axiophot.

Embryonic cuticle was examined by harvesting embryos from apple juice plates 24 hours after removing adults, dechorionating with bleach, and mounting in Hoyers/lactic acid. Slides were cleared by heating for several hours and then examined by dark field microscopy using a Zeiss Axiophot.

In situ hybridization was performed essentially as described (64), with probe prepared by nick translation of a double-stranded PCR product bearing the *bcd* ORF using alkali-stable digoxigenin-11-2’-deoxyuridine-5’-triphosphate (Roche), visualizing with alkaline-phosphate coupled sheep anti-digoxigenin Fab fragments (Roche) and subsequent reaction with 5-bromo-4-chloro-3-indolyl phosphate and nitro blue tetrazolium before examination using Nomarski optics on a Zeiss Axiophot.

For hand-sorting of embryos to identify those in nc 9-10, we followed the protocol of (65) with the following modifications. Embryos were dechorionated in 100% bleach for 1 minute, rinsed well, and then simultaneously devitellinized and fixed in a 1:1 ratio of ice-cold methanol and heptane for 5 minutes with gentle rocking. The vials were vortexed for 30 seconds, heptane and embryos at the interface removed, and the remaining embryos washed with ice-cold methanol 3 times. Embryos were then rehydrated into PBT with washing for 10 minutes and then stained with 0.1 μg/ml DAPI for 10 minutes. After washing in PBT, embryos were sorted under UV illumination using a Leica MZFLIII stereomicroscope.

### RNAseq and RT-qPCR

Total RNA was extracted from sorted embryos using TRIzol reagent (Thermo Fisher Scientific). Samples were submitted to the Ohio State University Cancer Center Genomics Shared Resource for poly(A) selection and library preparation with TruSeq (Illumina) and subsequently sequenced on a HiSeq4000 (Illumina) sequencer. Paired-end 150-nucleotide raw reads were first processed and demultiplexed at the sequencing facility, and subsequently processed as follows. Adapter sequences were trimmed followed by trimming low-quality bases from the 3’ end of reads based on a minimum Phred score of 15. Reads of less than 60nt were removed, as were reads mapping to mtRNA, tRNA, and rRNA. STAR (v2.5.3a) was used to map the reads either to the fly genome or the transcriptome (Dobin et al., 2013). Mapped reads were counted using RSEM (v1.3.0) (Li and Dewey, 2011). The resulting outputs were analyzed in R (v4.2.1) using the edgeR package (v3.40.2) to find differentially expressed genes and transcripts (Robinson et al., 2010). Low abundance genes and transcripts were filtered out by setting min.count=30 and min.total.count=30 in filterByExpr function in the edgeR package, removing genes and transcripts with < ∼2 counts per million mapped reads. All statistical analyses including differential expression and Locally Estimated Scatterplot Smoothing (LOESS) were performed using R (v4.2.1).

To account for over-sampling caused by the large depletion of mRNAs targeted by Upf1-Nos, correction factors were calculated as follows. We assumed that the most abundantly expressed mRNAs, which are transcribed from housekeeping genes, are not regulated by Upf1-Nos and thus can be used to calculate a correction factor. Rather than relying on a single highly expressed gene (e.g., eEF1α1), an average correction factor was calculated using the 20 most abundantly expressed genes for each Upf1-Nos replicate. Among genes with the highest average read abundance in wild type samples, we selected 20 with a similar rank order among Upf1-Nos and wild type samples. For each gene, the ratio of read abundance in the Upf1-Nos replicate to its average abundance across the three wild type replicates was calculated. The resulting ratios for all 20 genes were averaged, yielding a correction factor for the replicate (S2 Data). These correction factors were used as normalization factors for the Upf1-Nos replicates in edgeR. A similar approach was used to calculate three correction factors for the analysis of individual transcripts (S2 Data).

For RT-qPCR, cDNA was synthesized from 250 ng RNA using random hexamers and qPCR performed using SYBR Green using a 7500 Real Time PCR machine (Applied Biosystems). Triplicate measurements for each primer pair were obtained (for each of the three biological replicates). Abundance of each RT product was calculated relative to the abundance of RT product from RpS2 mRNA. Primer efficiencies for all primer pairs were 90%-100%.

### Motif enrichment analysis

Motif enrichment analysis was done using the following DREME command: “dreme -rna -norc -png -e 0.1 -m 10 -o DEOUT -p input.fa”. Parameters were set per the user manual. To prevent outlier large 3’-UTRs (e.g., the ∼10.5 kb *headcase* 3’-UTR) from biasing the analysis, only UTRs <2600 nt (98% of all 3’-UTRs) were analyzed.

### Gene ontology enrichment analysis

GO enrichment analysis was performed using DAVID (v6.8) with the default setting and categories (COG_ONTOLOGY, UP_KEYWORDS, UP_SEQ_FEATURE, GOTERM_BP_DIRECT, GOTERM_CC_DIRECT, GOTERM_MF_DIRECT, KEGG_PATHWAY, INTERPRO, SMART).

### Yeast interaction assays

Yeast were transformed with the 2µ plasmids described in Fig. 1 Supplement 1 by a standard lithium acetate/PEG protocol. For 2-hybrid experiments, we used the PJ69-4A strain (66), which is *MAT**a***, *trp1-901*, *leu2-3*, *112*, *ura3-52*, *his3-200*, *gal4*Δ, *gal80*Δ, *LYS2::GAL1-HIS3*, *GAL2-ADE2*, *met2::GAL7-lacZ*. Double transformants were obtained and grown on minimal SD dropout medium lacking tryptophan and leucine. For 3- and 4-hybrid experiments, we used the YBZ1 strain (67), which is *MAT**a**, ura3-52, leu2-3, 112, his3-200, trp1-1, ade2, LYS2::(LexAop)-HIS3, ura3::(lexA-op)-lacZ, LexA-MS2 coat (N55K)*. For 3-hybrid experiments, double transformants were grown on minimal SD dropout medium lacking uracil and leucine, and for 4-hybrid experiments triple transformants were grown on SD dropout medium lacking uracil, leucine, and tryptophan.

β-galactosidase activity was measured using Beta-Glo (Promega) essentially as described by Hook et al., 2005. Briefly, transformants were grown by diluting saturated overnight cultures into appropriate selective media to early-log phase (OD_600_ = 0.2 - 0.3), and then 40 µl of culture was incubated with the same volume of Beta-Glo reagent for 60 minutes at room temperature in 96-well microplates. Three transformants were grown separately and assayed for each experiment. Samples were analyzed in a Veritas luminometer (Turner Biosystems Inc). The output signal from the luminometer was divided by the OD_600_ to normalize for the number of cells in each sample, and by 1000 (by convention) to generate a reading of beta-galactosidase activity in arbitrary light units.

## Supporting information

Marhabaie Supplemental Data 1

Marhabaie Supplemental Data 2

Marhabaie Supplemental Data 3

Marhabaie Supplemental Data 4

Marhabaie Supplemental Data 5

Marhabaie Supplemental Data 6

## Acknowledgements

We thank Drs. K Panda, RK Slotkin, and R Bundschuch for advice and helpful discussions, Dr. P Yan for assistance with RNAseq sample preparation, Drs. G Singh and M Kearse for comments on the manuscript, H. Lipshitz for kindly forwarding their data for analysis in Fig. 5A, and Dr. C Croce for encouragement and support. We acknowledge services of the OSUCCC Genomics Shared Resource which is supported by P30CA016058. Some fly stocks were obtained from the Bloomington Drosophila Stock Center (NIH P40OD018537). This research was supported in part by NIGMS R01GM084376 (RPW).

## Supporting Information

**S1 Fig.**
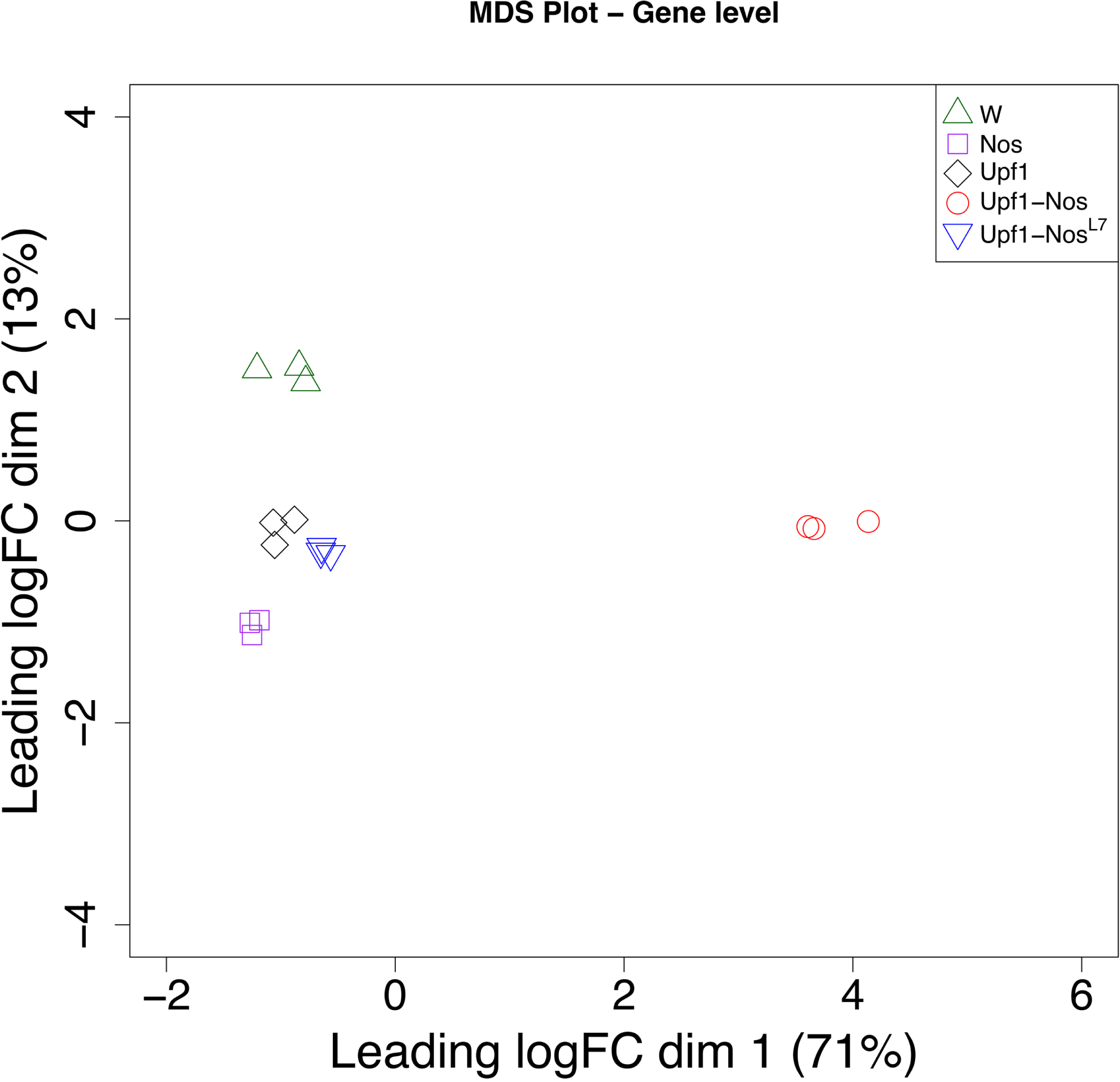
Multi-dimensional scaling plot of RNAseq data in Fig 2.

**S2 Fig.**
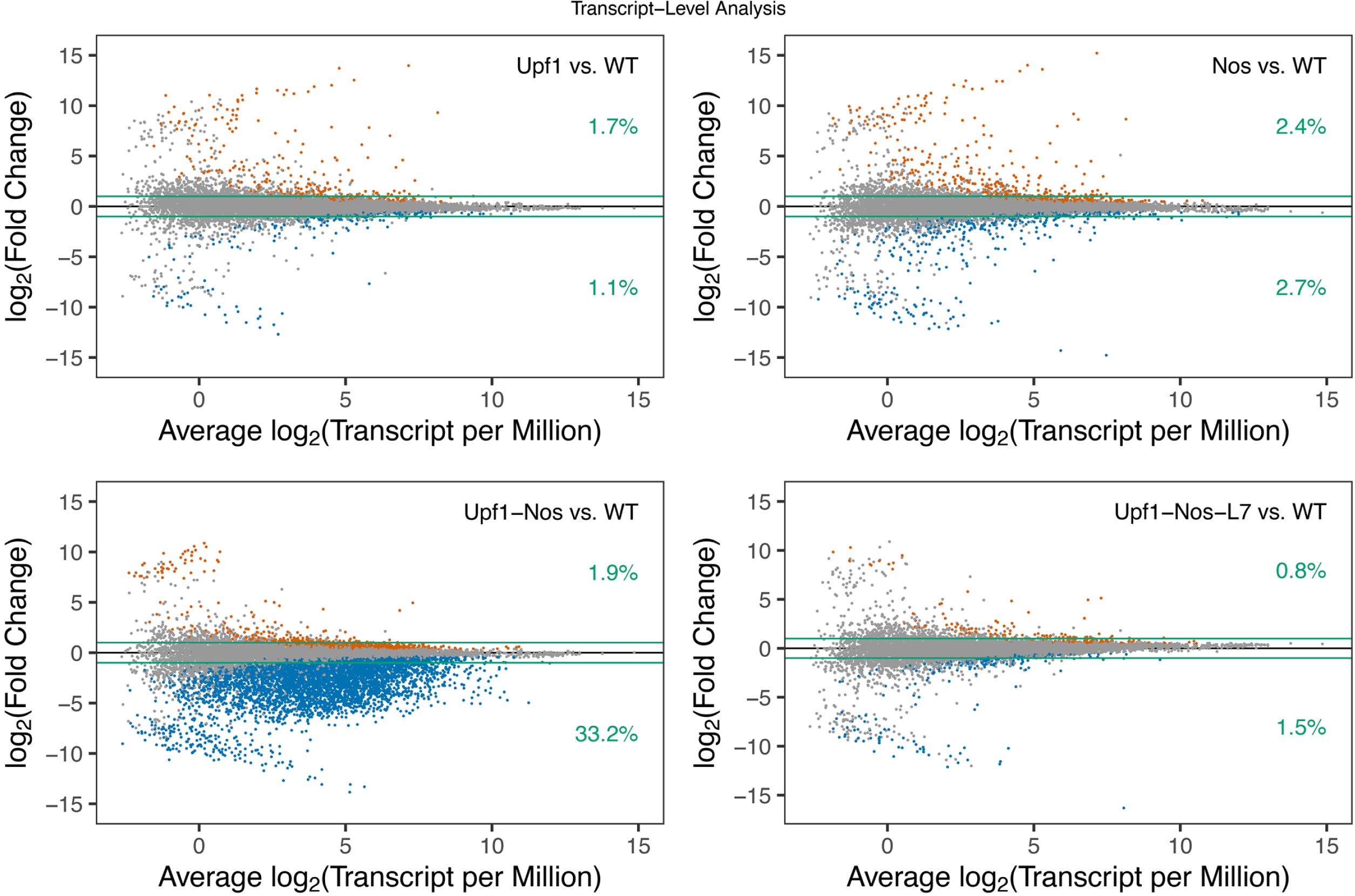
Smear plots of the RNAseq data plotted for individual mRNAs rather than gene-binned mRNAs (as in Fig 2A).

**S3 Fig.**
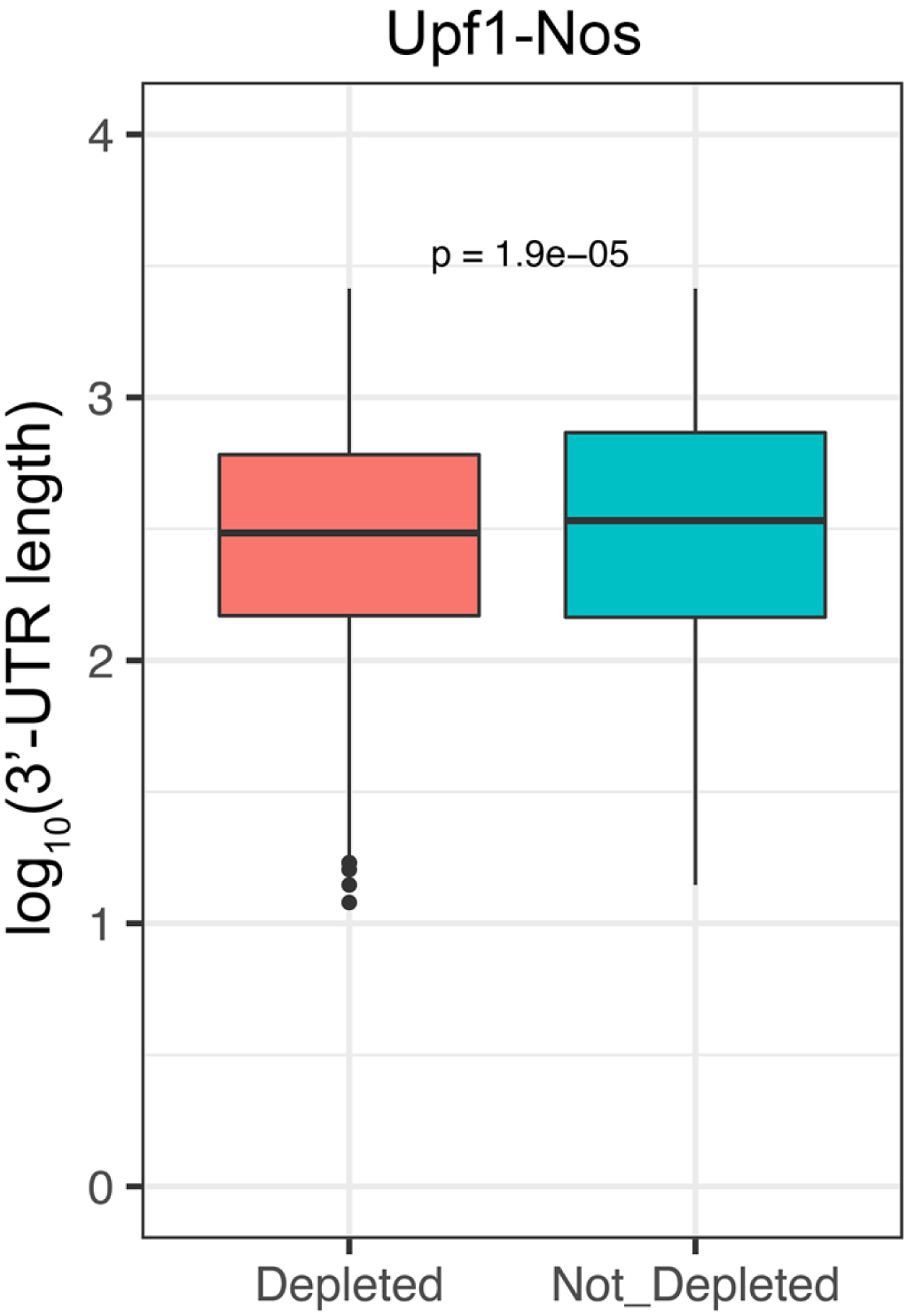
Boxplot showing that mRNAs targeted by Upf1-Nos have, on average, slightly shorter 3’-UTRs than non-targeted mRNAs.

**S4 Fig.**
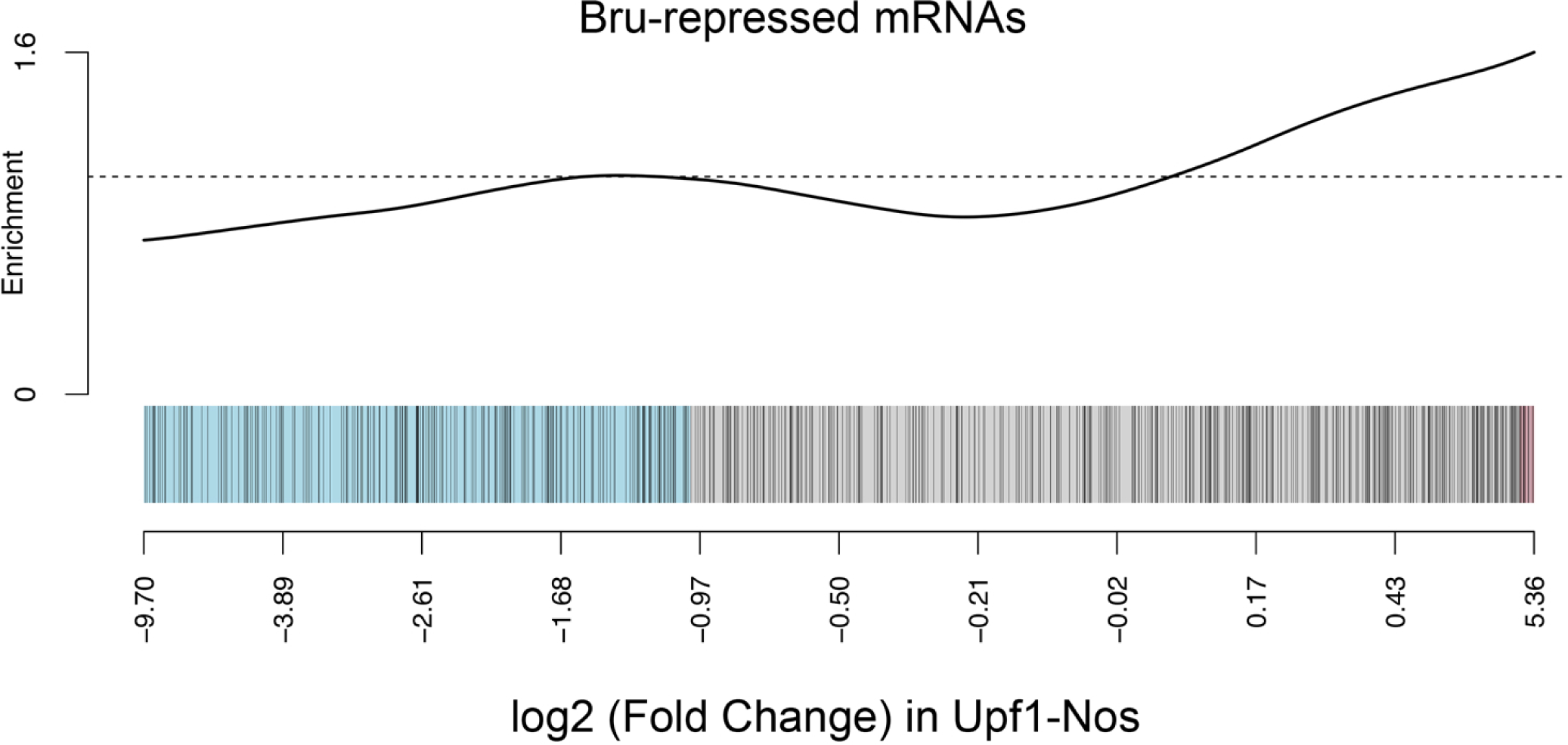
Barcode plot showing no significant enrichment of mRNAs targeted by Upf1-Nos among Bru-repressed mRNAs.

**S1 Data.** Data for Fig 1.

**S2 Data.** Data for Fig 2.

**S3 Data.** Data and statistical analysis for Fig 4.

**S4 Data.** Data and statistical analysis for Fig 5.

**S5 Data.** Data and statistical analysis for Fig 6.

**S6 Data.** Plasmids used in this work.

